# Multi-modal Proteomic Characterization of Lysosomal Function and Proteostasis in Progranulin-Deficient Neurons

**DOI:** 10.1101/2023.02.24.529955

**Authors:** Saadia Hasan, Michael S. Fernandopulle, Stewart W. Humble, Ashley M. Frankenfield, Haorong Li, Ryan Prestil, Kory R. Johnson, Brent J. Ryan, Richard Wade-Martins, Michael E. Ward, Ling Hao

## Abstract

Progranulin (PGRN) is a lysosomal protein implicated in various neurodegenerative diseases. Over 70 mutations discovered in the *GRN* gene all result in reduced expression of PGRN protein. However, the detailed molecular function of PGRN within lysosomes and the impact of PGRN deficiency on lysosomal biology remain unclear. Here we leveraged multifaceted proteomic techniques to comprehensively characterize how PGRN deficiency changes the molecular and functional landscape of neuronal lysosomes. Using lysosome proximity labeling and immuno-purification of intact lysosomes, we characterized lysosome compositions and interactomes in both human induced pluripotent stem cell (iPSC)-derived glutamatergic neurons (i^3^Neurons) and mouse brains. Using dynamic stable isotope labeling by amino acids in cell culture (dSILAC) proteomics, we measured global protein half-lives in i^3^Neurons for the first time and characterized the impact of progranulin deficiency on neuronal proteostasis. Together, this study indicated that PGRN loss impairs the lysosome’s degradative capacity with increased levels of v-ATPase subunits on the lysosome membrane, increased catabolic enzymes within the lysosome, elevated lysosomal pH, and pronounced alterations in neuron protein turnover. Collectively, these results suggested PGRN as a critical regulator of lysosomal pH and degradative capacity, which in turn influences global proteostasis in neurons. The multi-modal techniques developed here also provided useful data resources and tools to study the highly dynamic lysosome biology in neurons.

## Introduction

As the primary degradative organelle of the cell, the lysosome orchestrates proteostasis via the autophagy-lysosome pathway and degrades macromolecules such as proteins, lipids, carbohydrates, and RNA^1–3^. Neurons are particularly sensitive to lysosomal perturbations, as evidenced by numerous neurodegeneration-related mutations in genes that regulate lysosomal biology^4–6^. In particular, pathogenic mutations in genes that encode lysosomal or lysosome-associated proteins (e.g., *GRN*, *LRRK2*, *GBA*, *TMEM106B*, *C9*or*f72*) are major causes of inherited neurodegenerative diseases^5–7^. Genetic mutations associated with defective lysosomal enzymes lead to the accumulation of degradative substrates within the lysosomal lumen, consistent with chronic lysosomal dysfunction^8^. However, the molecular mechanisms by which many of these mutated genes cause lysosomal dysfunction and disease remain unclear.

Mutations in the *GRN* gene cause inherited frontotemporal dementia (FTD) and have also been linked to other neurodegenerative diseases, including neuronal ceroid lipofuscinosis (NCL), Alzheimer’s disease (AD) and Parkinson’s disease (PD)^9–12^. Over 70 pathogenic mutations in the *GRN* gene have been discovered, and all of these mutations result in reduced expression of progranulin (PGRN) protein^13–15^. Progranulin is trafficked to the lysosome and cleaved by cathepsins into smaller intra-lysosomal proteins called granulins^16^. Functionally, progranulin loss leads to a host of lysosome-related phenotypes, including defective autophagy and autophagosome-lysosome fusion^17,18^. Recently, lysosomal lipid dysregulation was found to be a major element of *GRN*-related disease pathogenesis^19,20^. However, the molecular cascade by which loss of intra-lysosomal progranulin impacts lysosomal biology and eventually leads to FTD remains elusive.

Capturing the dynamic lysosomal activities in highly polarized neurons is a challenging task, particularly in a high-throughput fashion. Our human induced pluripotent stem cells (iPSCs)-derived glutamatergic neuron (i^3^Neuron^21–23^) platform provides pure and scalable human neurons and can be genetically edited to create *GRN* null neurons as a neuronal model to study progranulin deficiency. Recent advances in capturing organelle dynamics have provided useful tools, such as proximity labeling in living cells via engineered ascorbate peroxidase (APEX^24^) or biotin ligases^25^, immunopurification of intact organelles^26^, and biotinylation by antibody recognition (BAR^27^) in primary tissues, though mostly in non-neuronal contexts. Other proteomics-based studies in progranulin mouse models mostly captured global changes regardless of cell type or organelle^28–30^. Developing proteomic techniques to probe lysosomes in neurons can provide valuable insights in the converging pathway of lysosomal dysfunctions in neurodegenerative diseases. We recently developed a lysosome proximity labeling method (Lyso-APEX) to characterize the dynamic lysosome interactome in wild-type (WT) i^3^Neurons^31–33^. In this study, we further expanded the lysosome toolbox by implementing the immunopurification of intact lysosomes (Lyso-IP) technique in our i^3^Neuron platform and Lyso-BAR technique in mouse brains. We comprehensively characterized lysosomal content and interactions using Lyso-APEX and Lyso-IP in i^3^Neurons and Lyso-BAR in fixed mouse brain tissues. To characterize global proteostasis in human neurons, we also designed a dynamic stable isotope labeling by amino acids in cell culture (dSILAC^34^) proteomic method that was suitable for iPSC-derived neuron cell type to measure global protein half-lives in i^3^Neurons for the first time.

Leveraging these multifaceted proteomic techniques, we systematically characterized the impact of progranulin loss using multi-modal readouts of lysosomal biology in i^3^Neurons and mouse brain. We found that loss of PGRN in human neurons presented increased levels of v-ATPase subunits on the lysosome membrane, increased catabolic enzymes within the lysosome, and elevated lysosomal pH. Mouse brains lacking PGRN also present elevated levels of lysosomal catabolic enzymes and bi-directional protein changes related to lysosomal transport. Using fluorescence microscopy, we confirmed that PGRN-deficient lysosomes are less acidic and have decreased cathepsin B activity compared to WT lysosomes. Consistent with impairments in protein homeostasis, *GRN* deficient i^3^Neurons have pronounced alterations in protein turnover, which was validated by FTD patient-derived i^3^Neurons carrying *GRN* mutation. Collectively, these results show that progranulin loss leads to a downstream molecular cascade involving lysosomal alkalinization and decreased degradative capacity, thereby impacting neuronal proteostasis. Multiple downstream proteins affected by these changes are involved in neurodegenerative pathways, suggesting molecular convergence of multiple neurodegeneration-related genes at the lysosome.

## Results

### Multi-modal proteomics captures holistic lysosomal biology

Lysosomes play critical roles in neurons such as degradation, endocytosis, signal transduction, nutrient sensing, and long-distance trafficking through axons^35–37^. Different methods of characterizing lysosomal composition and interactions now exist, each with its own strengths^33,38,39^. However, a comprehensive characterization of lysosomal biology in neurons with these modern tools has not been performed. We optimized and implemented three complementary proteomic strategies to characterize dynamic lysosomal interactions and lysosomal contents in both human neurons and mouse brain (**Figure 1A**). Lysosome proximity labeling using ascorbate peroxidase (Lyso-APEX) captured neuronal lysosome interactions with other cellular components as well as lysosome membrane proteins in living human neurons. Rapid lysosomal immunopurification (Lyso-IP) provided both lysosome lumen and membrane proteins in human neurons. Lysosomal biotinylation by antibody recognition (Lyso-BAR) revealed lysosome interactions *in situ* from fixed mouse brains. Proper location of these probes was validated by immunofluorescence and western blotting (**Figure 1B and Supplemental Figure S1**). Control groups were carefully selected for each probe to reduce nonspecific labeling and ensure intracellular spatial specificity (**Figure 1C**).

**Figure 1.**
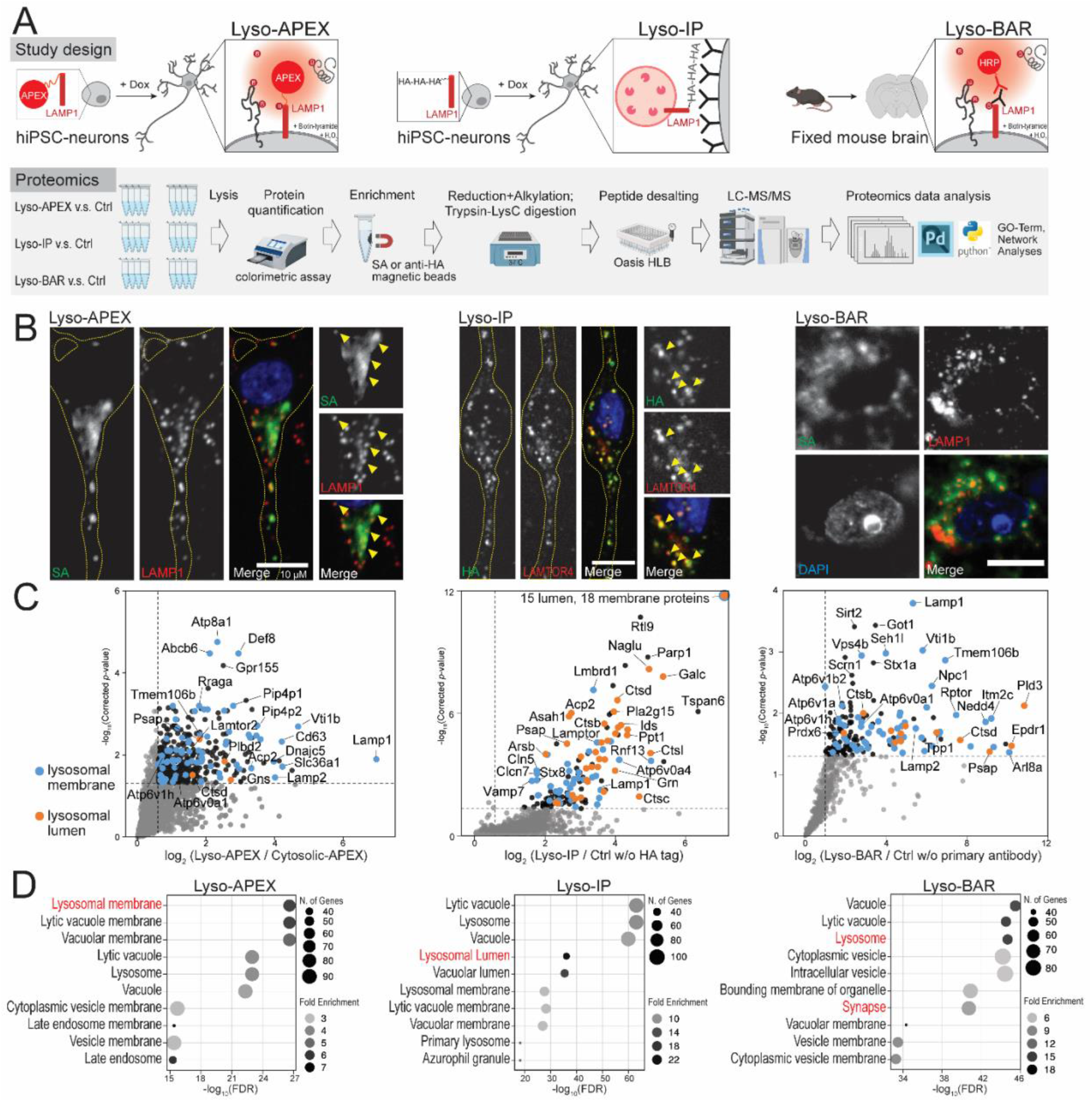
A map of the lysosomal proteome and interactome in human neurons and mouse brain. (**A**) Schematics of lysosomal proximity labeling (Lyso-APEX) in i^3^Neurons, lysosomal immunopurification (Lyso-IP) in i^3^Neurons, and lysosomal biotinylation by antibody recognition (Lyso-BAR) in fixed mouse brain tissues. (**B**) Fluorescence imaging of Lyso-APEX, Lyso-IP, and Lyso-BAR activities in i^3^Neurons and fixed mouse brain. Biotinylated proteins, stained with streptavidin (SA), colocalize with lysosomal markers in i^3^Neurons and fixed mouse brain tissues. HA-tagged lysosomes colocalize with lysosomal markers in i^3^Neurons. Scale bars are 10μm. (**C**) Volcano plots showing significantly enriched proteins from WT Lyso-APEX compared to cytosolic-APEX, Lyso-IP compared to control group without HA expression, and Lyso-BAR compared to control group without primary antibody staining (N=4). Dotted lines denote corrected *p*-value of 0.05 (y-axis) and ratio of 1.5 (x-axis). Known lysosomal membrane and lumen proteins are highlighted in blue and orange colors, respectively. (**D**) GO-term analyses of significantly enriched proteins in Lyso-APEX, Lyso-IP, and Lyso-BAR proteomics.

Lyso-APEX, Lyso-IP, and Lyso-BAR proteomics provided complementary coverage of the lysosomal microenvironment in human neurons and mouse brain tissues (**Figure 1D, Supplemental Figure S1**). Lysosomal membrane proteins such as v-ATPase subunits, LAMP proteins, and Ragulator subunits are identified and enriched by all three probes compared to corresponding controls. Lysosomal lumen proteins, especially hydrolases, are highly enriched in Lyso-IP proteomics, consistent with the degradative nature of the isolated organelles. Besides lysosome-resident proteins, both Lyso-APEX and Lyso-BAR proteomics captured dynamic lysosomal interaction partners related to organelle trafficking and axon transport (*e.g*., Kinesins, MAPs). Lyso-APEX favored surface-bound and surface-interacting proteins over luminal proteins due to the limited membrane permeability of reactive phenol-biotin generated on the cytosolic face of lysosomes during APEX-mediated labeling (**Figure 1C**). By contrast, Lyso-BAR revealed more intraluminal lysosomal proteins since BAR activation in fixed brain tissues requires membrane permeabilization. Lyso-BAR proteomics in mouse brain also captured numerous synaptic proteins, likely due to enhanced synaptic maturation *in vivo* compared to cultured iPSC-derived i^3^Neurons (**Figure 1D, Supplemental Figure S1**). Collectively, combining Lyso-APEX, Lyso-IP, and Lyso-BAR proteomic strategies allows us to obtain comprehensive lysosomal lumen and membrane proteomes as well as lysosomal interactomes in both cultured human i^3^Neurons and fixed mouse brains.

### Neuronal progranulin loss results in upregulation of vacuolar ATPases and alkalinization of lysosomal pH

Equipped with these new tools, we characterized how progranulin loss altered lysosomal biology. Using CRISPR-Cas9, we knocked out the *GRN* gene in wild type (WT) iPSCs harboring the Lyso-APEX probe and differentiated them into cortical neurons (**Figure 2A**). Immunofluorescence microscopy showed that progranulin protein colocalizes with lysosomes in WT i^3^Neurons as expected, and that no progranulin signal was observed in *GRN* KO i^3^Neurons (**Figure 2B**). Using Lyso-APEX proteomics, we found that PGRN depletion altered the abundance of many lysosome membrane proteins and lysosome interaction partners in human neurons (**Figure 2C)**. Gene Ontology (GO) enrichment analysis revealed upregulation of proteins related to lysosomal acidification and autophagy (**Figure 2D**). Numerous vacuolar ATPase (v-ATPase) and chloride channel proteins (CLCNs) were substantially up-regulated in *GRN* KO vs. WT i^3^Neurons (**Figure 2E, Supplementary Figure S2A**). GO enrichment analysis of significantly down-regulated proteins indicated impairment of lysosomal transport and RNA processing (**Supplementary Figure S2B**). Given the centrality of v-ATPases in establishing the acidic lysosomal lumen pH and the strong upregulation of acidification-related proteins in PGRN deficiency, we hypothesized that lysosomal pH could be perturbed by the loss of PGRN inside the neuronal lysosome^40^.

**Figure 2.**
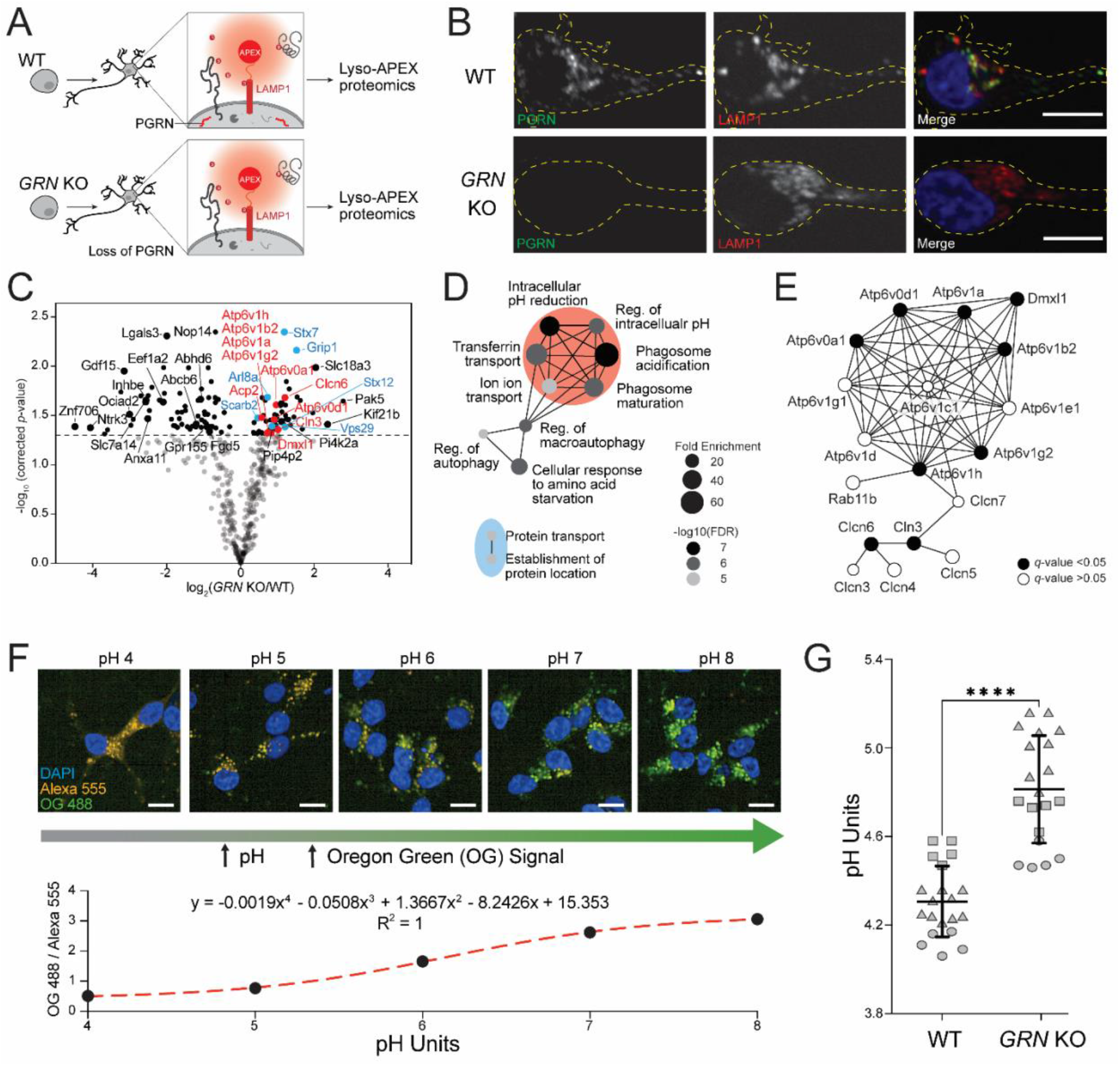
Lysosomal membrane proteins and pH are altered in i^3^Neurons with loss of progranulin. (**A**) Schematic of Lyso-APEX in WT and isogenic *GRN* KO i^3^Neurons. (**B**) Fluorescence imaging showing the colocalization of PGRN with lysosomes in WT i^3^Neurons and loss of progranulin (PGRN) signal in *GRN* KO i^3^Neurons. Scale bar is 10μm. (**C**) Volcano plot of Lyso-APEX proteomics in *GRN* KO vs. WT i^3^Neurons. Cytosolic enriched proteins and nonspecific labelings were removed from the volcano plot based on WT LysoAPEX vs. Cytosolic-APEX comparison. Red and blue colored proteins belong to lysosomal pH and protein transport GO-terms, respectively. (**D**) GO-term network analysis of significantly up-regulated biological processes in *GRN* KO vs. WT Lyso-APEX proteomics. (**E**) Protein network analysis of identified vacuolar-ATPase subunits and their interactors. (**F**) Live cell ratiometric lysosome pH assay. pH calibration curve is generated based on the ratio of pH-sensitive Oregon Green-488 dextran signal and pH-insensitive/loading control Alexa Fluor-555 red dextran in WT i^3^Neurons. Scale bar is 10μm. Other linear and nonlinear curve fitting models are provided in Supplementary Figure S2E. (**G**) Lysosome pH measurements in WT vs. *GRN* KO i^3^Neurons; multiple biological replicates from three independent experiments are represented with different shapes (**** denotes *p*-value < 0.0001).

To measure neuronal lysosomal acidification, we used a ratiometric fluorescent dextran assay. We co-generated an *in-situ* calibration curve using buffers of known pH, allowing accurate calculations of absolute pH within the lysosome with both nonlinear and linear curve fitting models (**Figure 2F, Supplementary Figure S2E**). Lysosomal pH is significantly increased in *GRN* KO i^3^Neurons (4.81 ± 0.24) compared to WT i^3^Neurons (4.31 ± 0.16). While this difference in pH may seem like a subtle change, it equates to a nearly three-fold decrease in the concentration of protons in the lysosomal compartment of *GRN* KO i^3^Neurons compared to WT counterparts due to the logarithmic nature of the pH scale ([H^+^] in WT ≈ 52±19 μM, *GRN* KO ≈ 18±9 μM). These observations show that *GRN* KO i^3^Neurons have alkalinized lysosomes, which could trigger the upregulation of acidification machinery to compensate for this effect.

### Progranulin-null lysosomes contain increased abundance of catabolic enzymes but have decreased enzymatic activity

Lysosomes require acidic luminal pH to degrade proteins using hydrolases^41^. Since lysosomes from progranulin-null neurons are less acidic, we hypothesized that these lysosomes may have altered abundances or activity of pH-dependent proteases. Using Lyso-IP proteomics, we characterized lysosome composition in *GRN* KO vs. WT i^3^Neurons (**Figure 3A**). PGRN protein was indeed enriched in WT Lyso-IP data and absent in *GRN* KO i^3^Neurons (**Supplementary Figure S3A**). Proteins involved in catabolism and lysosomal acidification were significantly increased in PGRN-deficient lysosomes in human neurons (**Figure 3B, 3C, Supplementary Figure S3B**). To investigate the impact of progranulin deficiency on lysosomes in mouse brain, we conducted Lyso-BAR proteomics in *GRN*^-/-^ vs. WT fixed mouse brains (**Figure 3D**). Similar protein catabolic processes were upregulated in *GRN*^-/-^ mice as indicated in Lyso-IP proteomics, particularly lysosomal proteases such as cathepsins (**Figure 3E, 3F, Supplementary Figure S3C**).

**Figure 3.**
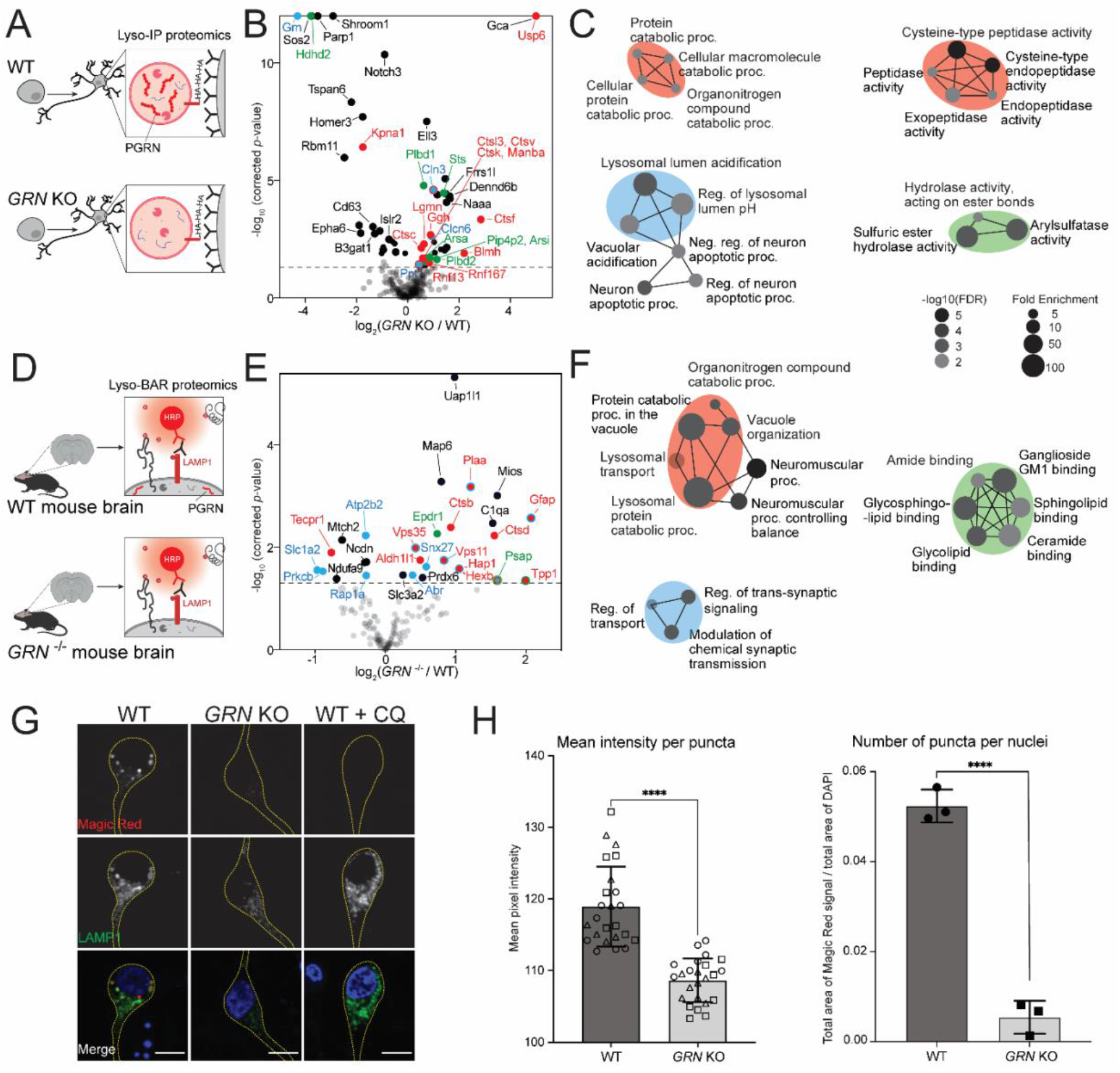
Progranulin-null lysosomes from human neurons and mouse brains have increased levels of lysosomal catabolic enzymes and decreased cathepsin B activity. (**A**) Schematic of intact lysosomal isolation (Lyso-IP) proteomics in *GRN* KO vs. WT i^3^Neurons. (**B**) Volcano plot of Lyso-IP proteomics showing protein changes related to protein catabolic processes (red), lysosomal pH (blue), and hydrolase activities (green). Nonspecific labeling proteins were removed based on WT LysoIP vs. control i^3^Neurons without HA expression. (**C**) GO-term network analysis of significantly changed proteins in *GRN* KO vs. WT Lyso-IP proteomics. Enriched biological processes are shown on the left. Molecular functions are shown on the right. Color code corresponds to the volcano plot. (**D**) Schematic of mouse brain Lyso-BAR labeling in *GRN*^-/-^ vs. WT mice. (**E**) Volcano plot showing Lyso-BAR protein changes in *GRN*^-/-^ vs. WT mouse brain. Nonspecific labeling proteins were removed based on WT LysoBAR vs. control without primary antibody staining. (**F**) GO-term network analysis of significantly changed proteins in *GRN* ^-/-^ vs. WT Lyso-BAR proteomics. (**G**) Fluorescence imaging of Magic Red assay to measure cathepsin B activity in i^3^Neurons. CQ stands for chloroquine treatment (50 μM for 24 hours). Scale bar is 10 μm. (**H**) Quantification of absolute and relative fluorescence intensities indicate decreased cathepsin B activity in *GRN* KO vs. WT i^3^Neurons (**** denotes *p*-value < 0.0001).

Prior studies of *GRN*^-/-^ mouse models have suggested that cathepsins may be less active in progranulin-null cells, despite increased abundance^30,42,43^. To directly evaluate the impact of progranulin depletion on lysosomal activity in human neurons, we quantified cathepsin B activity using a Magic Red assay in living WT and *GRN* KO i^3^Neurons. We observed a significant decrease in cathepsin B activity in PGRN-null i^3^Neurons compared to WT, indicating impaired proteolytic function (**Figure 3G, 3H**, **Supplementary Figure S3D**). To mimic alkalinization-related phenotypes observed in *GRN* KO i^3^Neurons, we treated neurons with chloroquine, an agent that neutralizes lysosomal pH. As predicted, direct alkalinization of lysosomes with chloroquine treatment reduced Magic Red fluorescence (**Figure 3G**). These findings confirm that although lysosomal hydrolases were upregulated in the absence of progranulin, their activity was decreased, likely due to alkalinized lysosomal lumens.

### Characterizing global protein turnover in human iPSC-derived neurons

Since lysosomes are major proteostatic organelles and their degradative function is impaired in progranulin-depleted neurons, we hypothesized that progranulin deficiency could influence global proteostasis. To measure the global protein turnover in neurons, we designed a dynamic SILAC proteomic method in cultured i^3^Neurons to obtain protein half-lives with multiple-time-point and single-time-point approaches (**Figure 4A**). By modeling the peptide degradation curves in WT i^3^Neurons, we found that most peptides’ degradation curves follow first-order exponential decay (**Figure 4B, Supplementary Figure S4A**). Peptide level and protein level half-lives correlate well with each other, with a median half-life of 4 days (**Figure 4C, 4D and Supplementary Figure S4B, S4C)**. Therefore, peptide and protein half-lives can be obtained using a single-time-point at 4 days (96 hrs) after heavy medium switch (**Supplementary Figure S4D)**. As we examined the distribution of protein half-lives, we found that numerous histones, nucleoporin proteins (Nups), proteins inside lysosomes as autophagy machinery (WDR45, GAA), and inner mitochondrial membrane proteins have extremely long half-lives (> 20 days) in i^3^Neurons, in agreement with recent studies in primary rodent neurons and brain tissues^44–46^. In contrast, proteins related to neurosecretion (GPM6B, VGF), axonal transport (kinesins), and ubiquitination (UBL4, USP11) have very short half-lives (0.3-2 days) (**Figure 4E, Supplemental Figure S4B**). Notably, one of the shortest half-life proteins in the entire neuronal proteome was STMN2, a microtubule-binding, golgi-localized protein implicated in ALS pathogenesis^47,48^. Lysosomal proteins have a median half-life of 3.6 days, slightly shorter than the median half-life of global neuronal proteins. Further investigation into the lysosomal compartment revealed a median half-life of 7.5 days for degradative enzymes, 3.5 days for V-ATPases, 6.2 days for lysosome-associated membrane glycoproteins (Lamps), 3.5 days for LAMTOR and HOPS complex subunits, and 3.1 days for BLOC1 complex subunits (**Figure 4F**). Together, this method enabled us to calculate global protein half-lives in live human i^3^Neurons for the first time.

**Figure 4.**
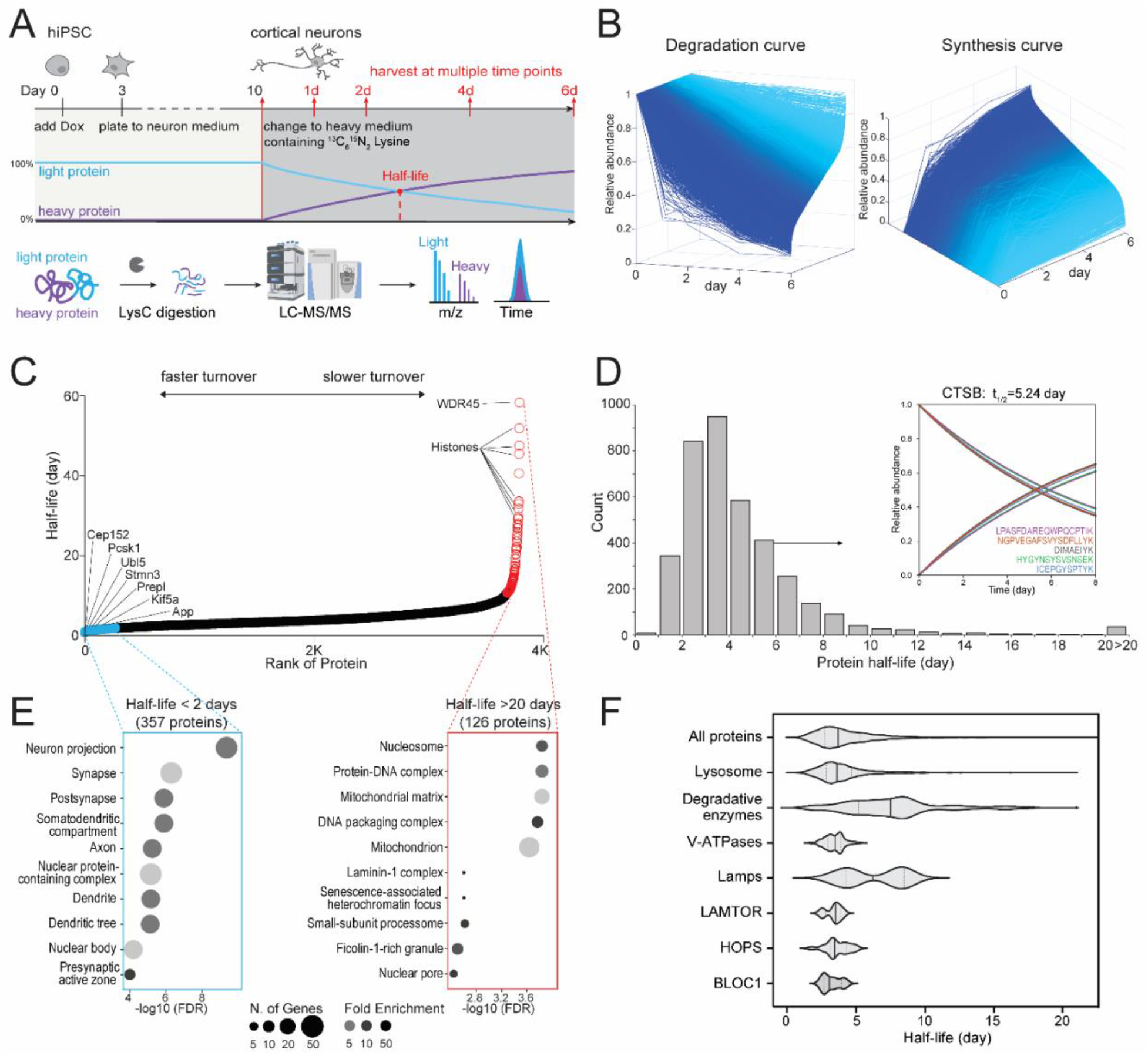
Measuring global protein half-lives in cultured human i^3^Neurons. (**A**) Schematic of dynamic stable isotope labeling by amino acids in cell culture (dSILAC) proteomics to measure global protein half-lives in cultured human i^3^Neurons. Cortical neurons were grown in normal medium until day 10 and then switched to heavy lysine-containing medium. Neurons are harvested at 1, 2, 4, and 6 days after medium switch followed by bottom-up proteomics. (**B**) Degradation and synthesis curves of all quantified proteins in WT i^3^Neurons. (**C**) Scatter plot of protein half-lives measured in WT i^3^Neurons ranked from fastest turnover to slowest turnover. (**D**) Histogram distribution of protein half-lives in WT i^3^Neurons. An example cathepsin B (CTSB) protein with five identified unique peptide sequences is illustrated in the inset. (**E**) GO-term analysis of the fast (left) and slow (right) turnover proteins in WT i^3^Neurons. (**F**) Violin plots of half-life distributions from all proteins and lysosomal proteins in WT i^3^Neurons.

### Progranulin deficiency alters neuronal protein turnover and decreases lysosomal degradative function

Using our dynamic SILAC proteomics approach, we evaluated protein turnover in WT vs. *GRN* KO i^3^Neurons (**Figure 5A**). The median of protein half-lives remained unchanged, but a remarkable 25% of all measured proteins presented significantly altered half-lives in *GRN* KO vs. WT i^3^Neurons (**Figure 5B and 5C**). Proteins related to polymerization and fiber organization showed significantly slower turnover, which may indicate a propensity for protein misfolding and aggregation in *GRN* KO neurons related to FTD pathogenesis (**Figure 5D**)^49^. Despite the significantly slower turnover of both cathepsin B and cathepsin D, proteins related to RNA catabolic processes showed faster turnover, which further implicates the disturbance of molecular degradation pathways (**Figure 5E**). Several proteins with altered half-lives (either faster or slower turnover) are involved in ALS/FTD and other neurodegenerative diseases, suggesting potential converging pathways among different neurodegenerative diseases and dysfunction of key regulators of proteostasis (**Figure 5F**).

**Figure 5.**
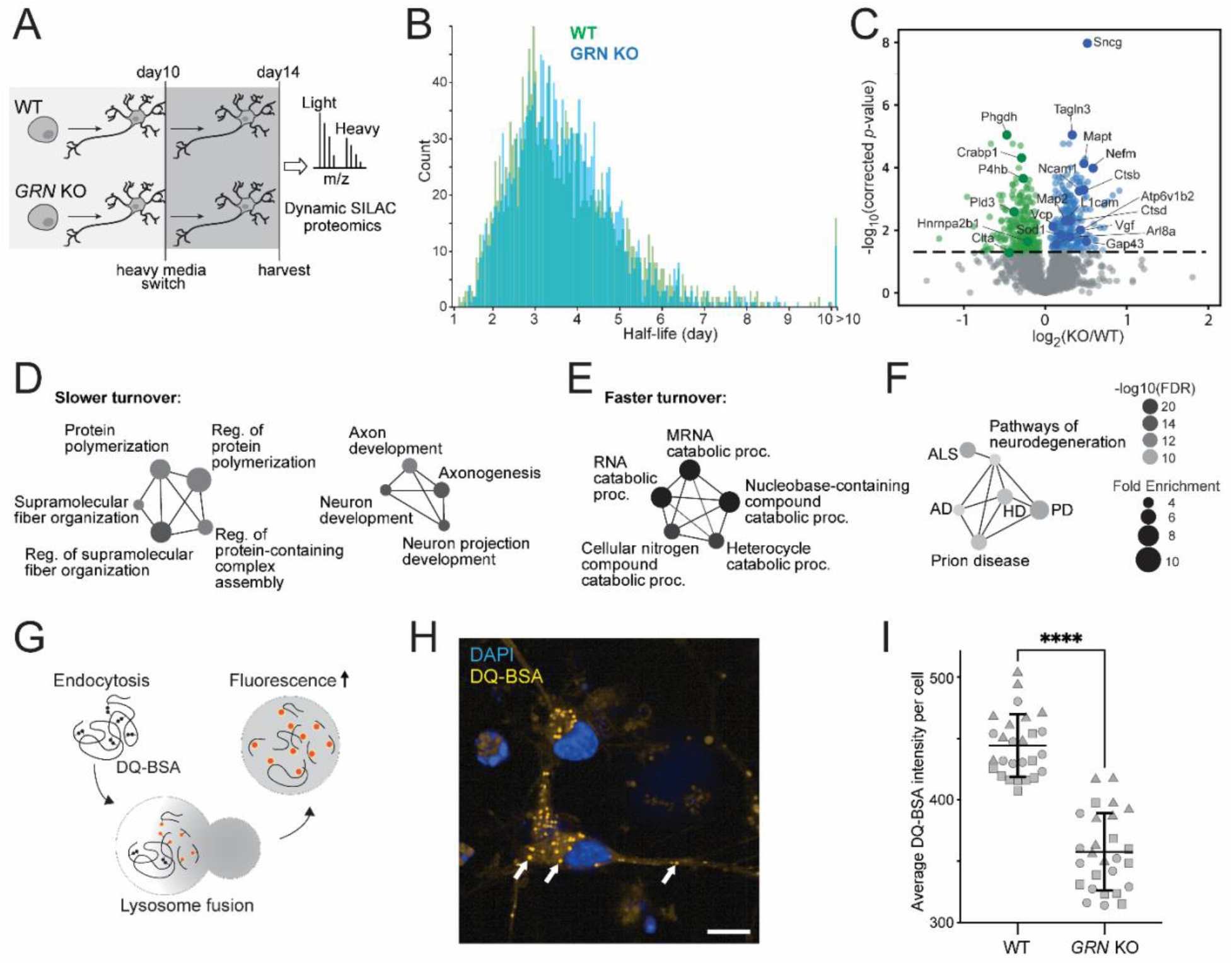
Global protein turnover and lysosomal degradative function are impaired in i^3^Neurons with loss of progranulin. (**A**) Schematic of protein half-life measurements in *GRN* KO vs. WT i^3^Neurons using dynamic SILAC proteomics. (**B**) Histogram distribution of global protein half-lives in *GRN* KO (blue) vs. WT (green) i^3^Neurons. (**C**) Volcano plot of protein half-life changes in *GRN* KO vs. WT i^3^Neurons. (**D**) GO-term network analysis of enriched biological processes from proteins with significantly slower turnover. (**E**) GO-term network analysis of enriched biological processes from proteins with significantly faster turnover. (**F**) KEGG pathways enriched from significantly altered protein half-lives in *GRN* KO vs. WT i^3^Neurons. (**G**) Schematic of the DQ-BSA Red assay to measure lysosomal degradative function. Extracellular DQ-BSA with self-quenched dye is endocytosed into i^3^Neurons and trafficed to the lysosome, where it isdegraded into smaller protein fragments with isolated fluorophores with fluorescence signals. (**H**) Representative fluorescence imaging of DQ-BSA Red assay showing DQ-positive lysosomes in i^3^Neurons. Scale bar is 10μm. (**I**) Quantification of the fluorescence intensities of the DQ-BSA Red assay in WT vs. *GRN* KO i^3^Neurons, normalized to the total number of puncta in two groups (**** denotes *p*-value < 0.0001).

Given our observations that lysosomes within *GRN* KO i^3^Neurons are alkalinized, have reduced cathepsin activity, and exhibit major changes in global protein homeostasis, we predicted that *GRN* KO lysosomes would exhibit impaired lysosome-mediated protein degradation. We directly assayed lysosomal degradative capacity using a fluorescent DQ-BSA Red assay (**Figure 5G, 5H**)^50,51^. The DQ-BSA substrate is initially self-quenching due to the close spatial proximity of the fluorophores. Once cleaved in acidic lysosomes, the DQ-BSA substrate exhibit bright fluorescence signals. The mean DQ-BSA intensity in *GRN* KO i^3^Neurons was significantly decreased compared to WT neurons (**Figure 5I**), similar to pharmacological inhibition of lysosomal degradation using chloroquine (**Supplemental Figure S5A**). The change in active proteolysis was independent of lysosomal biogenesis, as there was no change in the number of puncta per cell in *GRN* KO vs. WT (**Supplemental Figure S5B**). Taken together, these results show that *GRN* KO lysosomes have significantly hindered proteolytic capacity, consistent with our observations of pathological impairment in lysosomal acidification and impaired lysosomal hydrolase activity.

### An isogenic series of *GRN* mutation patient-derived iPSC neurons exhibit altered protein homeostasis

To further explore the relationship between *GRN* insufficiency and protein homeostasis abnormalities, we created i^3^Neurons from an FTD patient-derived iPSC line with a heterozygous *GRN* mutation^52^ (c.26 C>A, p.A9D; referred subsequently as ptMut), as well as the isogenic iPSC control line with corrected *GRN* mutation (ptWT). We further knocked out *GRN* in this control line to create an additional isogenic *GRN* KO iPSC line (ptKO) (**Figure 6A**). After differentiating each of these lines to i^3^Neurons, performing dSILAC, and measuring their protein half-lives, we found that over 25% of proteins showed significantly altered half-lives in ptKO compared to ptWT i^3^Neurons (**Figure 6B**), consistent with *GRN*-KO vs. WT comparison in **Figure 5C**. About 15% of protein half-lives were significantly altered in ptMut compared to ptWT group (**Figure 6B and 6C**). Principal component analysis and hierarchical clustering showed complete separations of both genetic background and *GRN* genotypes from five i^3^Neurons lines (*GRN*-KO, WT, ptKO, ptMut, ptWT) based on protein half-lives (**Figure 6D, 6E**). The overall protein half-life changes also suggested a potential gene dosage effect, in which many proteins have greater fold changes in *GRN*-KO neurons compared to *GRN*-mutant neurons (**Figure 6F, Supplementary Figure S6A**). Half-life changes of key overlapping proteins in the three comparisons (*GRN*-KO vs. WT, ptKO vs. ptWT, ptMut vs. ptWT) are highlighted in **Figure 6G and Supplementary Figure S6B**.

**Figure 6.**
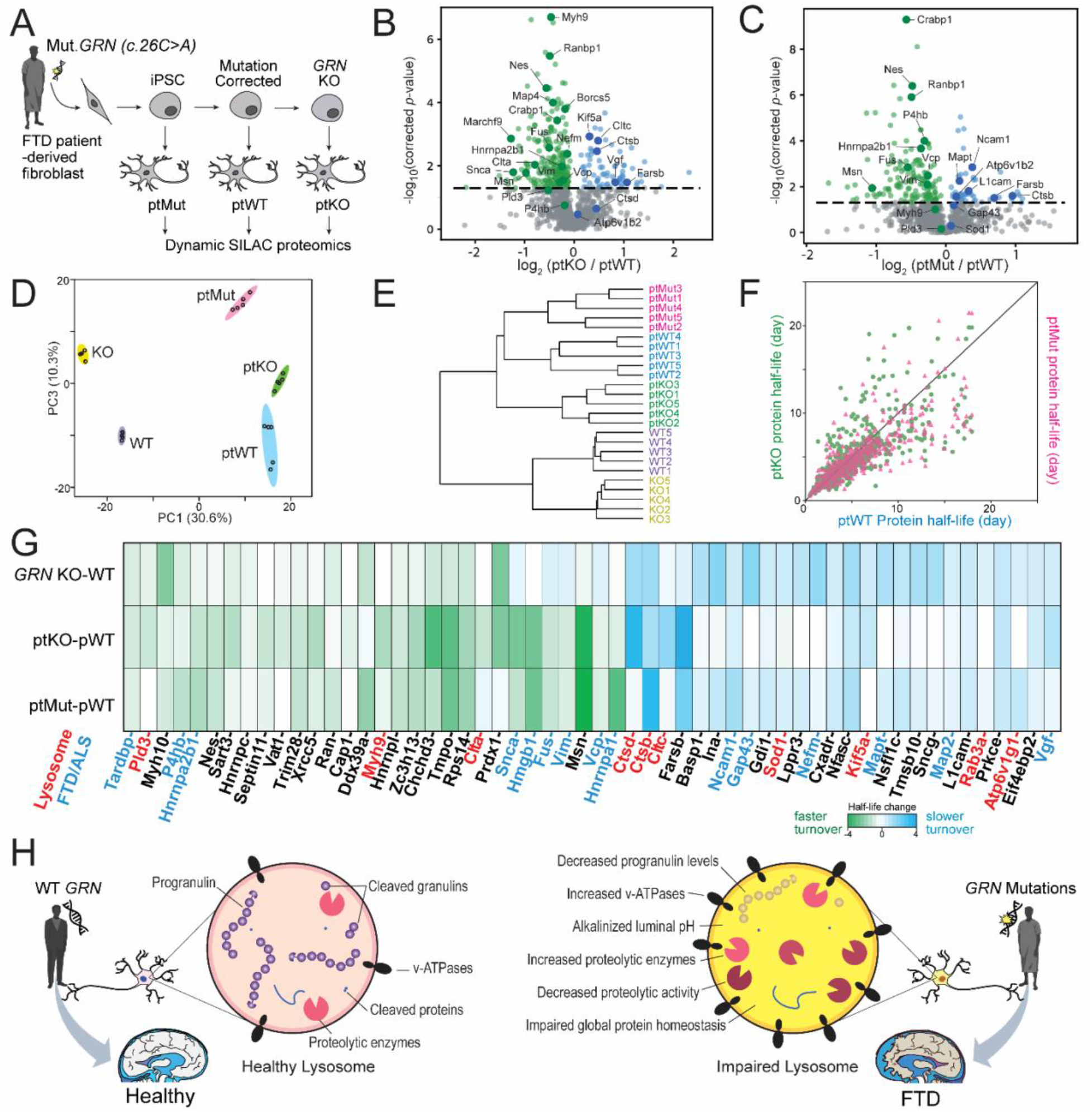
Frontotemporal dementia (FTD) patient-derived i^3^Neurons with muatant *GRN* reveal altered protein turnover of lysosomal enzymes and FTD-associated proteins. (**A**) Generation of a set of FTD patient fibroblast-derived i^3^Neurons. First, CRISPR-Cas9 was used to insert an inducible *NGN2* cassette into the *AAVS1* locus of a patient fibroblast-derived iPSC line (ptMut). Next, CRISPR-Cas9 was used to correct the *GRN* mutation in ptMut to create an isogenic control iPSC line (ptWT) and then to knockout *GRN* in pWT to create the ptKO iPSC line. These iPSC lines were then differentiated into i^3^Neurons and dSILAC proteomics was performed. (**B**) Volcano plot of protein half-life changes in ptKO vs. ptWT i^3^Neuron. (**C**) Volcano plot of protein half-life changes in ptMut vs. ptWT i^3^Neuron. (**D**) Principal component analysis using protein half-lives in GRN-KO, WT, ptKO, ptMut, and ptWT i^3^Neurons groups. (**E**) Hierarchical clutering of five i^3^Neurons groups. (**F**) Scatter plot of protein half-life changes in ptKO vs.ptWT and ptMut vs. ptWT comparisons showing the consistency and potential gene dosage effect of ptKO and ptMut i^3^Neurons. (**G**) Heatmap showing key overlapping protein turnover changes in *GRN* KO vs. WT, ptKO vs. ptWT, and ptMut vs. ptWT i^3^Neurons. Heatmap colors represent the absolute half-life differences between comparison groups. Key proteins from lysosomes and relevant to FTD/ALS are highlighted in red and blue, respectively. (**H**) Schematic of proposed lysosomal impairment in progranulin-deficient neurons caused by GRN mutations in FTD patients.

The findings in patient-derived *GRN* mutant and KO neurons validate our observations of dysregulated protein homeostasis in settings of *GRN* depletion and insufficiency, including alterations in the half-lives of numerous neurodegeneration-associated proteins. Many lysosomal enzymes showed prolonged protein half-lives, such as cathepsins (CTSD, CTSB), which was especially notable given our direct measurements of increased cathepsin levels and reduced CTSB activity in *GRN* KO neurons. Our findings additionally show that substantial upregulation of numerous lysosomal-associated proteins and enzymes occurs in *GRN*-deficient neurons – many via prolongation in protein half-lives – but that these homeostatic changes are insufficient to normalize lysosomal degradative capacity. As summarized in **Figure 6H**, we propose that GRN mutations that cause PGRN deficiency inside neuronal lysosomes result in alkalinized lysosomal pH, decreased proteolytic activities, and impaired global protein homeostasis that eventually lead to frontotemporal dementia.

## Discussion

Lysosomal dysfunction is a convergent pathological mechanism across multiple neurodegenerative diseases^5,6^. Progranulin, a glycoprotein linked to FTD, ALS, PD, and AD, is trafficked to, processed by, and resides within the lysosome^15^. Despite this knowledge, the primary molecular functions of progranulin and the impact of progranulin deficiency on lysosomal biology and protein homeostasis remain unclear. This is in part due to limited tools available for understanding the role of progranulin in the highly dynamic lysosomes in the brain. Here, we designed a combination of *in vitro* and *in situ* proximity labeling, lysosome immunopurification, and dynamic SILAC proteomic approaches to map the organellar and cellular architectures of neuronal progranulin deficiency.

For the first time, we implemented the antibody-guided biotinylation strategy to measure lysosomal composition in the brain and the lysosomal immunopurification method to characterize neuronal lysosomes. We additionally developed and optimized a neuron dynamic SILAC proteomic method to calculate protein half-lives in i^3^Neurons for the first time. Despite the application of dynamic SILAC in various cell culture and mouse models, it remains challenging to measure protein turnover rates in non-dividing cells, particularly in human neurons^44,45,53^. Many neuronal proteins exhibit extremely long half-lives, particularly nuclear proteins due to a lack of cell division. For the first time, we measured the global protein turnover in i^3^Neurons and found that the dynamics of most proteins can be modeled using first-order exponential decay. This enabled the measurement of global neuron protein half-lives using a 4-day single time point method, significantly reducing the starting materials and reagents compared to multiple-time-point method and allowing the streamlined comparison of multiple i^3^Neuron lines with different genome backgrounds and *GRN* genotypes.

Using these new multi-modal proteomic strategies, we discovered that progranulin deficiency leads to increased expression of v-ATPases on the lysosomal membrane in i^3^Neurons. Upon further investigation, we discovered that progranulin deficiency had a severe impact on the lysosomes’ ability to properly acidify, which results in impaired hydrolytic activity despite an upregulation of acidification machinery. These results suggest that progranulin plays an important role in maintaining lysosomal pH, with v-ATPases either contributing to that effect or providing a compensatory response for that effect. Since alkalinized lysosomes cannot properly hydrolyze substrates, we next looked at how the contents of progranulin-null lysosomes were affected. We found that several lysosomal enzymes were upregulated both in the mouse and human dataset, notably cathepsins. We showed decreased cathepsin B activity in live neurons, a phenomenon only shown in *in vitro* assays before^52,54–56^. Similar perturbations of lysosomal acidification have been reported in non-neuron cells and other neurodegenerative diseases^57,58^.

Mutations in the *GRN* gene cause progranulin deficiency inside the lysosome and have been shown to impair lysosomal function and the autophagy pathway^13,18^. However, whether progranulin deficiency alters protein turnover in human neurons has not been systematically investigated previously. We found that progranulin deficiency broadly influenced proteostasis, altering the half-lives of over 15% and 25% neuron proteins in *GRN* mutant and KO i^3^Neurons, respectively. Lysosomes degradative capacity was compromised by PGRN deficiency, as evidenced in our DQ-BSA assay. Critically, the recapitulation of global proteostasis defects in FTD-patient-derived neurons suggests that altered protein turnover rates are relevant to disease pathophysiology.

Although we have established exciting new tools and characterized the neuronal lysosome quite extensively, there are several limitations in this study. Although LAMP1 is a classic lysosome marker, it is also expressed on late endosomes and other endocytic species^59^. Despite this limitation, our data is consistent with degradative lysosome proteomics, and we obtained new insights into neuronal lysosomes specifically. We also recognize that human iPSC-derived neurons are not fully mature and representative of late-stage disease, and therefore have supplemented i^3^Neuron data with lysosomal proteomics in aged mice. As neuron is the major cell type of the brain, LysoBAR proteomics provide consistent and complementary lysosomal changes compared to cultured i^3^Neuron. However, LysoBAR method is not cell-type specific and will also include lysosome profiles from other cell types such as microglia, which has higher expression level of progranulin compared to neurons^9,15^. It will be important to investigate whether aged human neurons exhibit different proteomic changes and if human microglial lysosomes behave differently compared to neurons in future studies. Furthermore, future research can focus on individual proteins with altered lysosomal enrichment and half-lives as novel handles for elucidating disease mechanisms, discovering disease biomarkers, and further assessing whether these neuronal proteostatic changes manifest in established mechanisms of neurodegenerative pathology, such as stress granule persistence, impaired macroautophagy, and failed fusion of lysosomes to autophagosomes.

Overall, this study developed and implemented a set of novel proteomics techniques to decipher neuronal lysosomal biology and proteostasis in the context of *GRN* insufficiency that causes frontotemporal dementia. We provided new insights of progranulin function in regulating lysosomal pH, lysosomal catabolic activity, and global proteostasis in neurons, opens numerous avenues for future follow-up studies to determine specific molecular mechanisms underpinning the protein changes discovered here. This work also illustrated a roadmap for how multi-modal proteomics can be used to illuminate lysosomal biology, providing useful data and technical resources that can be applied to characterize other organelle dynamics in neurons.

## Methods

### Human i^3^Neuron Culture

Human iPSC-derived cortical neurons (i^3^Neurons) were cultured based on our previously established protocol^23^. Briefly, human iPSCs were maintained on Matrigel (Corning Incorporated #354277) coated tissue culture dishes in Essential 8 medium (Gibco #A1517001). Several iPSC lines were used in this study as listed in **Table 1**. A doxycycline-inducible neurogenin2 (NGN2) cassette (Addgene #105840) was stably integrated into each iPSC line, enabling rapid differentiation to glutamatergic cortical neurons (i^3^Neurons) in a week. Between day 0 and day 3, iPSCs were maintained in neuronal induction medium^23^. Day-3 neurons were replated on poly-L-ornithine coated plates in Brainphys neuron medium and maintained by half-medium change every two days until neuronal maturation in two weeks.

**Table 1:**
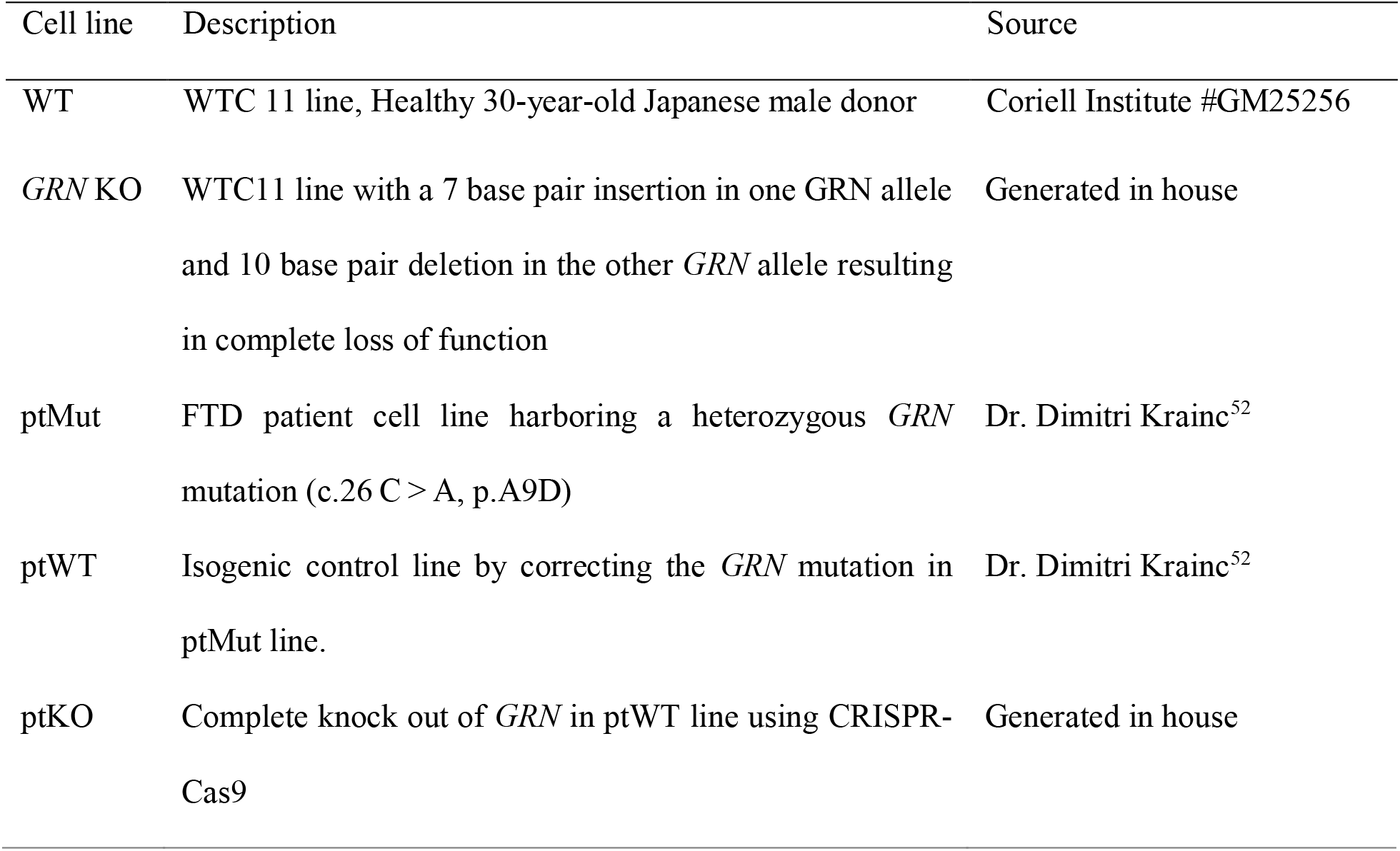
List of human iPSC lines used in this study.

| Cell line | Description | Source |
| --- | --- | --- |
| WT | WTC 11 line, Healthy 30-year-old Japanese male donor | Coriell Institute #GM25256 |
| *GRN* KO | WTC11 line with a 7 base pair insertion in one *GRN* allele and 10 base pair deletion in the other *GRN* allele resulting in complete loss of function | Generated in house |
| ptMut | FTD patient cell line harboring a heterozygous *GRN* mutation (c.26 C > A, p.A9D) | Dr. Dimitri Krainc<sup>52</sup> |
| ptWT | Isogenic control line by correcting the *GRN* mutation in ptMut line. | Dr. Dimitri Krainc<sup>52</sup> |
| ptKO | Complete knock out of *GRN* in ptWT line using CRISPR-Cas9 | Generated in house |

### Animals

All mice used in this study were obtained from the Jackson Laboratory and housed in the NIH animal facility that followed NINDS/NIDCD/NCCAM Animal Care and Use Committee (ACUC) Policy for animal husbandry and euthanasia. WT (C57BL/6J) and *GRN*^-/-^ (B6.129S4(FVB)-Grntm1.1Far/Mmjax, MMRRC stock#036771-JAX) mice were used here^60^. Whole brains were dissected from 20-month-old male WT and *GRN*^-/-^ mice after cardiac perfusion with 4% paraformaldehyde (PFA). Cortex was fixed in 4% PFA overnight, incubated in 30% sucrose for 24 hours, and snap frozen on dry ice. A microtome was used to generate 40 μm thick coronal slices that were stored in cryoprotectant at −30°C.

### Lysosomal proximity labeling in i^3^Neurons

Lysosomal proximity labeling was achieved by stable integration of ascorbate peroxidase (APEX2) enzyme onto the C terminus of LAMP1 protein in human iPSCs and differentiating iPSCs into i^3^Neurons, as the previously established KuD-LAMP1-APEX (Lyso-APEX) line^32^. A cytosolic localized nuclear exporting signal (NES) APEX i^3^Neuron line was used as the spatial control^31,32^. Prior to proximity labeling, i^3^Neurons were incubated in 500 μM biotin-tyramide (Adipogen, #41994-02-9) for 30 min in a CO_2_ incubator. Proximity labeling was induced by incubating the cells in 1 mM of hydrogen peroxide for exactly 1 min followed by rapid quenching using ice-cold quench buffer (10 mM sodium azide, 10 mM sodium ascorbate, 5 mM TROLOX in PBS). Neurons were lysed with cold lysis buffer (50 mM Tris-Cl pH 7.4, 500 mM NaCl, 0.2% SDS, 1 mM DTT, 10 mM sodium <u>azide</u>, 10 mM sodium ascorbate, 5 mM TROLOX, cOmplete mini protease inhibitor tablets). Detailed sample preparation procedures have been described previously^33^. Briefly, neuron lysates were sonicated with QSonica (Q800R) sonicator for 15 min at 2°C and clarified by centrifugation. Total protein concentrations were measured using a detergent-compatible (DC) Colorimetric Protein Assay (Bio-Rad #5000111). Biotinylated proteins were enriched with streptavidin (SA) magnetic beads (Cytiva, # 28-9857-99) for 18 h rotating at 4°C and washed extensively to reduce non-specific bindings. Biotinylated proteins were reduced, alkalized, and digested into peptides on the SA beads. The optimal SA beads-to-protein ratio and trypsin-to-SA beads ratio were previously determined^32^. After overnight digestion using Trypsin/Lys-C (Promega, #V5073), supernatant was collected from the magnetic beads, and the digestion reaction was quenched with 10% trifluoroacetic acid until pH < 3. Peptides were desalted with a Waters Oasis HLB 96-well extraction plate, dried under SpeedVac, and stored at −30 °C until LC-MS analysis.

### Rapid Lysosome Immunopurification from i^3^Neurons

Lysosome Immunopurification (Lyso-IP) iPSC line was generated by the stable expression of LAMP1-3xHA in WT and *GRN* KO iPSC lines. i^3^Neurons were differentiated as described above and maintained in 15cm dishes until day 14. A control i^3^Neurons line without HA expression (mEmerald) was used to control nonspecific labeling background. Neurons were washed 2 times with PBS and dissociated from the plate using forceful pipetting of 10 ml of PBS. Next, neurons were resuspended in 1ml cold KPBS (136 mM KCl, 10 mM KH2PO4, pH 7.25 adjusted with KOH) and gently homogenized with 21 strokes through an isobiotec balch-style cell homogenizer with a 10μm ball bearing. Each neuron lysate sample was incubated with 150 μL of pre-washed anti-HA magnetic beads (Thermo #88836/88837) for 3 min on a rotator and gently washed three times with KPBS. Beads bound with intact lysosomes were resuspended in 100 μl of Lyso-IP lysis buffer (50 mM HEPES, 50 mM NaCl, 5 mM EDTA, 1% SDS, 1% TritonX, 1% NP-40, 1% Tween 20, 1% deoxycholate, 1% glycerol, 5 mM TCEP) and heated at 60 °C for 30 mins at 1000 g agitation. The supernatant was collected, and the beads were washed with an additional 50 μl of lysis buffer. Supernatant was combined into a new tube for routine bottom-up proteomics steps as described below.

### Lysosomal proximity labeling in mouse brains

Mouse brain slices were picked evenly throughout the whole brain and washed with PBS three times. Endogenous peroxidase activity in brain slices was quenched with 0.3% H_2_O_2_ in PBS for 30 minutes. The slices were blocked using 3% donkey serum and 0.25% Triton X in PBS followed by primary antibodies in blocking buffer at 4°C on a rocker overnight. After the slices were washed thoroughly with PBST, they were incubated with secondary antibody conjugated to HRP in blocking buffer for 1 hour at room temperature and extensively washed in PBST. The slices were then incubated in biotin-tyramide with 1% fetal bovine serum (FBS) in PBS for 30 min, and then 0.003% H2O2 for 10 min, immediately followed by washing with quench buffer (10 mM sodium azide and 500 mM sodium ascorbate). Brain slices without primary antibody treatment were used as the negative control group to compare with Lyso-BAR. One slice from each group was further treated with appropriate Alexa Fluor for microscopy imaging. Twenty brain slices from each group were transferred to 100 μL of Lyso-BAR lysis buffer (3% SDS + 2% sodium deoxycholate in PBS), boiled at 99°C for 1 hour at 1200 g agitation, and sonicated with QSonica Sonicator for 15 min. The lysate was boiled again at 99°C for an additional 30 min until all tissues were homogenized and dissolved into solution. The lysate was diluted using PBS to reduce SDS concentration and clarified by centrifugation. Biotinylated proteins were enriched following the same steps described above for Lyso-APEX sample preparation with optimized SA beads-to-protein ratio and trypsin-to-beads ratio for Lyso-BAR samples.

### Dynamic SILAC proteomics in i^3^Neurons

Human i^3^Neurons were maintained on PLO coated 12-well dishes in light amino acid-containing media (DMEM:F12 for SILAC medium (Athena Enzyme Systems #0423), N2 Supplement (Life Technologies Corporation #17502048), B27 Supplement (Life Technologies #NC1001496), NEAA (Life Technologies #11140050), GlutaMAX (Life Technologies #35050061), BDNF (PeproTech #450-02), NT-3 (PeproTech #AF-450-03-100ug), 0.3 mM of Arginine (Sigma #A4599), and 0.5 mM of Lysine (Sigma #L7039)). On day 10 of i^3^Neuron culture, neurons were gently washed with PBS twice and switched into media containing the same components except for replacing light lysine with heavy stable isotope labeled (^13^C6^15^N_2_) lysine (Cambridge Isotope Laboratories #CNLM-291-H-PK). For multiple time point experiments, neurons were harvested at 1, 2, 4, and 6 days (accurate to within 10 min) after media switch. For single time point experiments, neurons were harvested after 4 days (96 hours) of media switch. Neurons were gently washed with PBS twice, lysed in 100 μL of ice-cold lysis buffer containing 0.1% Rapigest (Waters #186008740), 150 mM NaCl, and 50 mM Tris-HCl, sonicated for 15 min, and clarified by centrifugation. Total protein concentrations were determined by DC Protein assay (BioRad). Protein disulfide bonds were reduced by 5 mM of Tris(2-carboxyethyl) phosphine (TCEP) for 30 min, followed by addition of 15 mM of iodoacetamide (IAA) for 30 min in a ThermoMixer shaking at 800 g at 37°C. Proteins were digested with LysC (Promega #VA1170) at 1:30 (enzyme:protein) ratio for 16 hours at 37°C and quenched with 10% trifluoroacetic acid (TFA) until pH<3. Peptides were desalted using a Waters Oasis HLB 96-well extraction plate based on the manufacturer’s protocol. Peptide samples were dried under SpeedVac and stored at −80°C until LC-MS analysis.

### LC-MS/MS analysis

LC-MS/MS analyses were conducted on a Dionex UltiMate3000 nanoLC system coupled with a Thermo Scientific Q-Exactive HFX or a Fusion Lumos mass spectrometer. Before injection, peptide samples were reconstituted in 2% acetonitrile (ACN), 0.1% formic acid (FA) in LC-MS grade water and centrifuged to collect supernatant. Easy-spray PepMap C18 columns (2 μm, 100 Å, 75 μm ×75 cm) were used for peptide separation with a flow rate of 0.2 μL/min and column temperature of 60°C. The mobile phase buffer A was 0.1% FA in water, and buffer B was 0.1% FA in acetonitrile. A two-hour gradient was used for proximity labeling proteomics, and a three-hour gradient was used for SILAC proteomics. LC-MS/MS analyses were conducted with a top 40 data dependent acquisition with MS range of m/z 400-1500, MS resolution of 120K, isolation window of m/z 1.4, dynamic exclusion of 22.5 s, and collision energy of 30% for higher-energy collisional dissociation (HCD) fragmentation. Automatic gain control (AGC) targets were 1×10^6^ for MS and 2×10^5^ for MS/MS. Maximum injection times were 30 ms for MS and 35 ms for MS/MS.

### Proteomics Data analysis

LC-MS/MS raw files from Lyso-APEX, Lyso-IP, and Lyso-BAR proteomic experiments were analyzed with Thermo Fisher Proteome Discoverer (2.4.1.15) software. For dynamic SILAC proteomic data, MaxQuant (1.6.17.0) software was used for data analysis. Swiss-Prot *Homo sapiens* database was used for i^3^Neuron data and *Mus musculus* database was used for mouse data with 1% false discovery rate (FDR) for protein identification. Custom-made contaminant protein libraries (https://github.com/HaoGroup-ProtContLib) were included in the data analysis pipeline to identify and remove contaminant proteins^61^. Trypsin was selected as the enzyme with a maximum of two missed cleavages. Cysteine carbamidomethylation was included as fixed modification, and oxidation of methionine and acetylation of the protein N-terminus were selected as variable modifications.

Protein/peptide identification and peak intensities were output as excel files for further analysis using Python or R. Statistical analyses (t-test) and volcano plots for Lyso-APEX, Lyso-IP, and Lyso-BAR proteomics were conducted in Python. Lyso-APEX and Lyso-BAR data were normalized to the most abundant endogenously biotinylated protein (PCCA) before statistical analysis as described previously^32^. For dynamic SILAC data, Maxquant output files were further processed with R to calculate heavy/light peptide ratios and construct the degradation and synthesis curves as well as curve-fitting to the first-order kinetic in multiple time point experiment. For single time point experiments, peptide level Maxquant output files were processed with Python to calculate the peptide half-lives using the equation: t_1/2_ = t_s_ × [ln2 / ln (1+Ψ)], where t_s_ represents the sampling time after media switch, and Ψ represents the heavy-to-light abundance ratio of the peptide. Protein level half-lives were calculated by averaging the half-lives of unique peptides belonging to the specific protein. Statistical analysis was conducted with t-test, and multiple half-life datasets were merged by uniprot protein accession in Python. Protein GO enrichment analysis was conducted using ShinyGO^62^. Protein network analysis was conducted with STRING^63^.

### Live Cell Ratiometric pH Assay

Live cell ratiometric lysosomal pH measurements were conducted using a modified method from Saric et al^64^, further optimized for high content imaging and analysis. WT and *GRN*-KO i^3^Neurons were maintained on 96-well dishes. On day 10, neurons were loaded with 50 μg/mL pH-sensitive Oregon Green-488 dextran (Invitrogen, #D7171), and 50 μg/mL pH-insensitive/loading control Alexa Fluor-555 red dextran (Invitrogen, #D34679) for 4 hours, before washing three times with PBS then chased overnight with neuronal media after PBS washes the day before imaging. These dextrans accumulate in lysosomes, and high-content microscopy quantification of their fluorescence enables ratiometric calculations of pH within individual lysosomes. Physiological buffers of known pH (4-8) containing 10 μg/mL nigericin were placed on WT neurons to generate a calibration curve. Live cell spinning disk confocal microscopy was performed using a Opera Phenix HCS System (PerkinElmer); calibration and sample wells were imaged at 63×; counterstaining was done with NucBlue Live ReadyProbes Reagent (Invitrogen, #R37605) to count and segment nuclei. Lysosome pH was calculated as ratiometric measurement of lysosomes (488/555nm), with subsequent calculation of the pH of those compartments based on the corresponding calibration curve. All analysis was performed using PerkinElmer’s Harmony HCA Software (PerkinElmer). Statistical analyses for all imaging data were conducted using independent student’s t-test.

### Magic Red cathepsin B activity assay

Human i^3^Neurons were plated at a density of 50,000 cells on PLO-coated ibidi slides (Ibidi # 80827) and maintained to day 10. Magic Red (Abcam #AB270772-25TEST) was added to the cells at 1:25 final dilution and incubated in the dark for 30 mins at 37°C. Cells were washed twice with PBS and incubated with Hoechst 33342 (Thermo Scientific #62249) at 1:10,000 for 5-10 mins and then washed with PBS. Neurons were imaged using Nikon spinning disk confocal at 60× oil objective. Images were edited and analyzed using ImageJ software^65^. Statistical analysis was conducted using independent student’s t-test.

### Live cell DQ-BSA Assay in i^3^Neurons

WT and *GRN*-KO i^3^Neurons were plated on 384-well dishes. On day 10, neurons were incubated with 45 μg/mL DQ-BSA Red (Invitrogen, #D12051) for 5 hours to allow for substrate endocytosis. Live-cell spinning disk confocal microscopy was performed via Opera Phenix HCS System (PerkinElmer); control and sample wells were imaged at 40× and counterstaining was done with NucBlue Live ReadyProbes Reagent (Invitrogen, #R37605) to count and segment nuclei. All analysis was performed via PerkinElmer’s Harmony HCA Software (PerkinElmer).

### Western blotting

Intact lysosomes were isolated via immunopurification as described above. The intact lysosomes on beads were boiled with sample buffer at 95 °C for 5 mins. The beads were magnetized, and the supernatant was used for immunoblotting. Lysates were separated using 4-15% precast polyacrylamide gels (Bio-Rad, # 4561083) at **100 V** and then transferred using the Trans-Blot Turbo transfer kit onto nitrocellulose membranes (Bio-Rad, #1704270). Membranes were blocked with 5% nonfat dry milk prepared in TBST (Tris-buffered saline with Tween 20) for 1 hour at room temperature and probed with primary antibodies in 5% bovine serum albumin (BSA) in TBST at 4°C overnight (See **Table 2** for antibodies and dilutions). Following incubation, membranes were washed 3× with TBST and incubated in secondary antibodies diluted 1:5000 in 5% BSA for 1 hour at room temperature. Membranes were then washed 3× with TBST and visualized using ECL western blotting substrate.

**Table 2.** Antibodies used for immunostaining.

| Antibody | Company | Catalog | Application | Dilution |
| --- | --- | --- | --- | --- |
| LAMP1 | Developmental Studies Hybridoma Bank | #H4A3 | Immunofluorescence | 1:3000 |
| LAMTOR4 | Cell Signaling Technology | #13140S | Immunofluorescence | 1:500 |
| HA | Millipore Sigma | #11867423001 | Immunofluorescence | 1:500 |
| HOECHST | Millipore Sigma | #63493 | Immunofluorescence | 1:10000 |
| Streptavidin-680 | Jackson ImmunoResearch | #016-620-084 | Immunofluorescence | 1:1000 |
| Streptavidin-488 | Jackson ImmunoResearch | #016-540-084 | Immunofluorescence | 1:1000 |
| PGRN | R&D systems | #AF2420 | Immunofluorescence | 1:1000 |
| Rhodamine Red Donkey anti-Mouse IgG (H+L) | Jackson ImmunoResearch | #715-295-151 | Immunofluorescence | 1:1000 |
| Rhodamine Red Donkey anti-Rabbit IgG (H+L) | Jackson ImmunoResearch | (#711-295-152 | Immunofluorescence | 1:1000 |
| Rhodamine Red Donkey anti-Rat IgG (H+L) | Jackson ImmunoResearch | #711-295-153 | Immunofluorescence | 1:1000 |
| Alexa Fluor 488 Donkey anti-Mouse IgG | Jackson ImmunoResearch | #715-545-151 | Immunofluorescence | 1:1000 |
| Rodamine Red Donkey anti-Goat IgG | Jackson ImmunoResearch | #705-295-147 | Immunofluorescence | 1:1000 |
| LAMP2 | Cell Signaling Technology | #D5C2P | Western blot | 1:5000 |
| Calreticulin | Cell Signaling Technology | #D3E6 | Western blot | 1:5000 |
| CTSD | Proteintech | #21327-I-AP | Western blot | 1:5000 |
| Catalase | Proteintech | #2126-1-AP | Western blot | 1:5000 |
| P70 S6 Kinase | Cell Signaling Technology | #49D7 | Western blot | 1:5000 |
| PGRN | R&D systems | #AF2420 |  |  |

### Fluorescence imaging

i^3^Neurons were cultured on PLO-coated ibidi slides (Ibidi #81506) for fluorescence imaging. Neurons were fixed in 4% PFA for 10 mins, washed very gently with PBS, and incubated in blocking buffer (1% bovine serum albumin + 0.1% TritonX) for 1 hour at room temperature (RT). Next, neurons were incubated with primary antibody in blocking buffer overnight at 4°C, gently washed with PBS, and incubated in secondary antibody for 1 hour at RT. Following thorough washes, neurons were ready to be imaged. Mouse brain slices were prepared in the same steps as neuron culture for fluorescence imaging. All antibodies and their respective applications and dilutions are listed in **Table 2**. Confocal images were obtained using a Nikon Eclipse Ti spinning disk confocal microscope at 60× using an oil immersion objective with constant setting between experimental groups. Data analysis was conducted in ImageJ.

## Author Contributions

Author contributions: L.H. and M.E.W. initiated the project with help from S.H. and M.S.F. to design the experimental plan. S.H., M.S.F., S.W.H., R.P., and L.H. conducted iPSC-neuron culture and sample preparation. S.H. conducted mouse brain sample preparation. S.H., M.S.F., A.M.F., H.L., and L.H. conducted proteomics experiments. A.M.F., K.J., and L.H. performed proteomics data analysis. S.H. and M.S.F. performed western blotting and microscopy experiments. S.W.H., S.H., and B.J.R. conducted live cell assays. S.H. and L.H. wrote the manuscript with input from M.S.F. and M.E.W. and edits from all coauthors.

The authors declare no competing financial interests.

## Acknowledgements

This study is supported by the NIH grant (R01NS121608, Hao), GW intramural grant (UFF, Hao), the Intramural Research Program at NIH/NINDS (Ward), and MRC Dementias Platform UK Stem Cell Network Capital Equipment Awards MR/M024962/1 (Wade-Martins). We would like to thank Maia Parsadanian for her assistance with mouse brain immunostaining and cell culture, Dr. Dimitri Krainc for providing the FTD patient iPSC lines, and Dr. Lingnan Lin for his help in Python data analysis.

## Data Availability

All proteomics RAW files have been deposited in the PRIDE database (ProteomeXchange Consortium) with the data identifier PXD040251 and will be released upon publication. All other supporting data are available within the article and the supplementary files.

