## Supplementary for "Multi-modal Proteomic Characterization of Lysosomal Function and Proteostasis in Progranulin-Deficient Neurons"

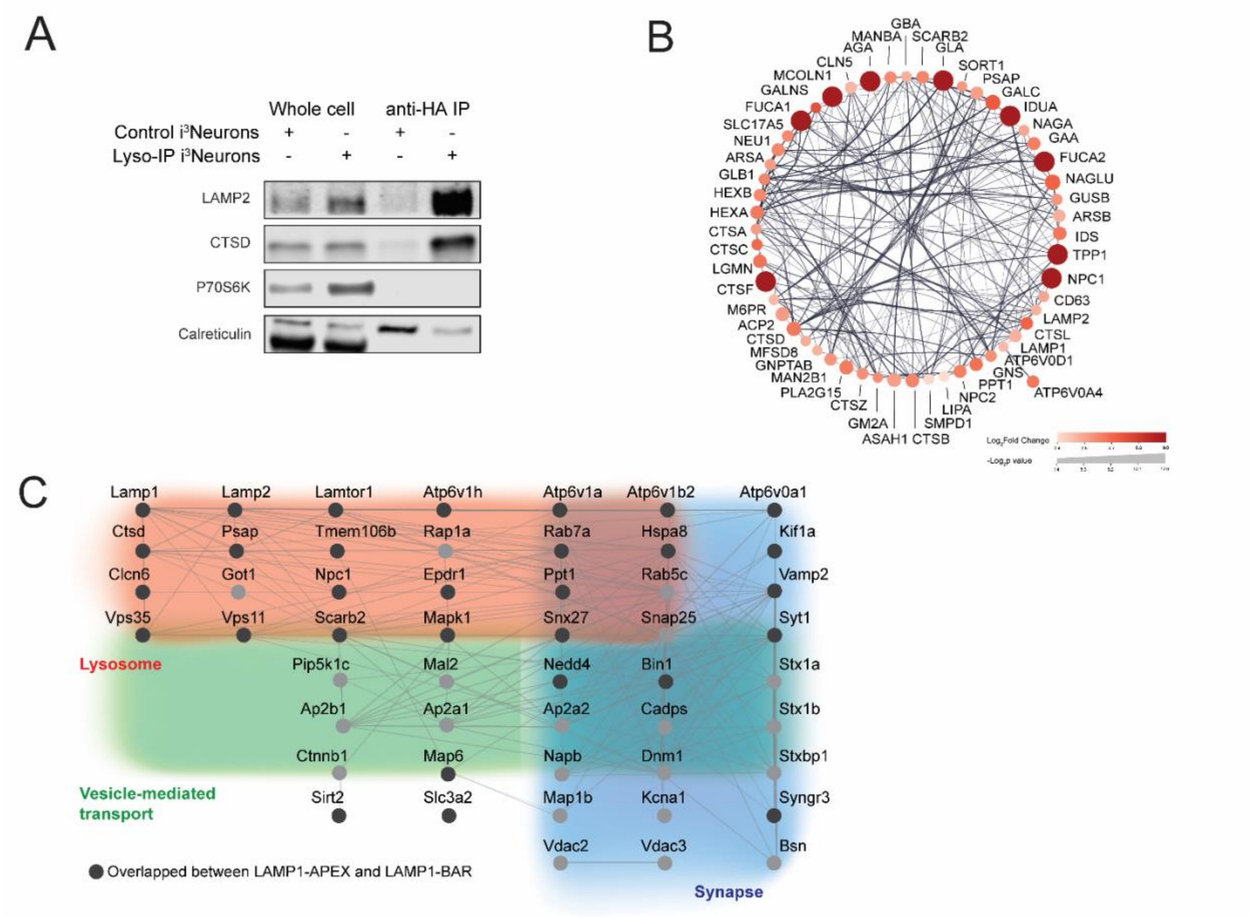


**Supplementary Figure S1. Confirmation of the lysosomal locations of Lyso-IP, Lyso-BAR, and Lyso-APEX probes, related to Figure 1.** (**A**) Western blot analysis of isolated lysosomes from Lyso-IP i$^3$Neurons. (**B**) Protein network analysis of enriched lysosomal proteins from Lyso-IP neurons. (**C**) Protein network analysis of proteins enriched in Lyso-Bar mouse brain samples, highlighting the proteins overlapping with the Lyso-APEX probe in cultured iPSC-derived neurons.

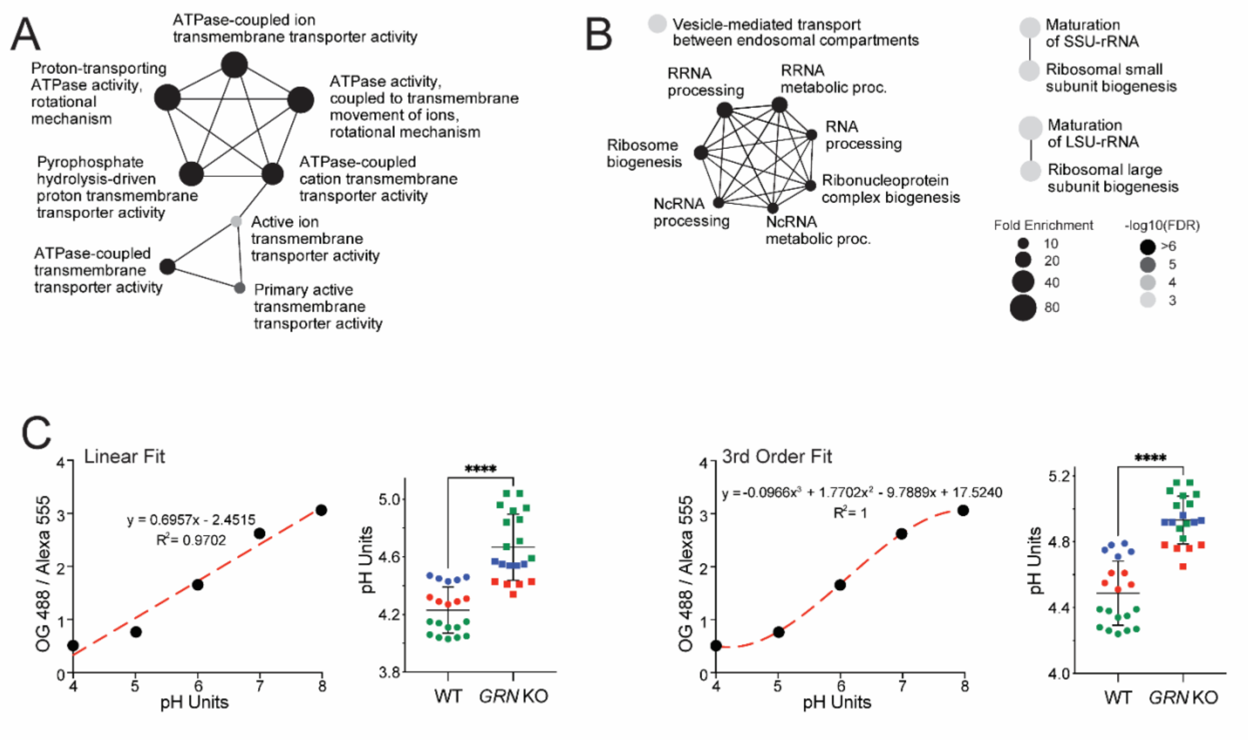


**Supplementary Figure S2. Proximity labeling proteomics and lysosome pH measurement in i³Neurons with loss of progranulin, related to Figure 2. (A)** GO-term network analysis of significantly up-regulated molecular functions in *GRN* KO vs. WT Lyso-APEX proteomics. **(B)** GO-term network analysis of significantly down-regulated molecular functions in *GRN* KO vs. WT Lyso-APEX proteomics. **(C)** Lysosomal pH measurements in WT vs. *GRN* KO i³Neurons with linear (left) and 3rd order (right) calibration curve fitting.

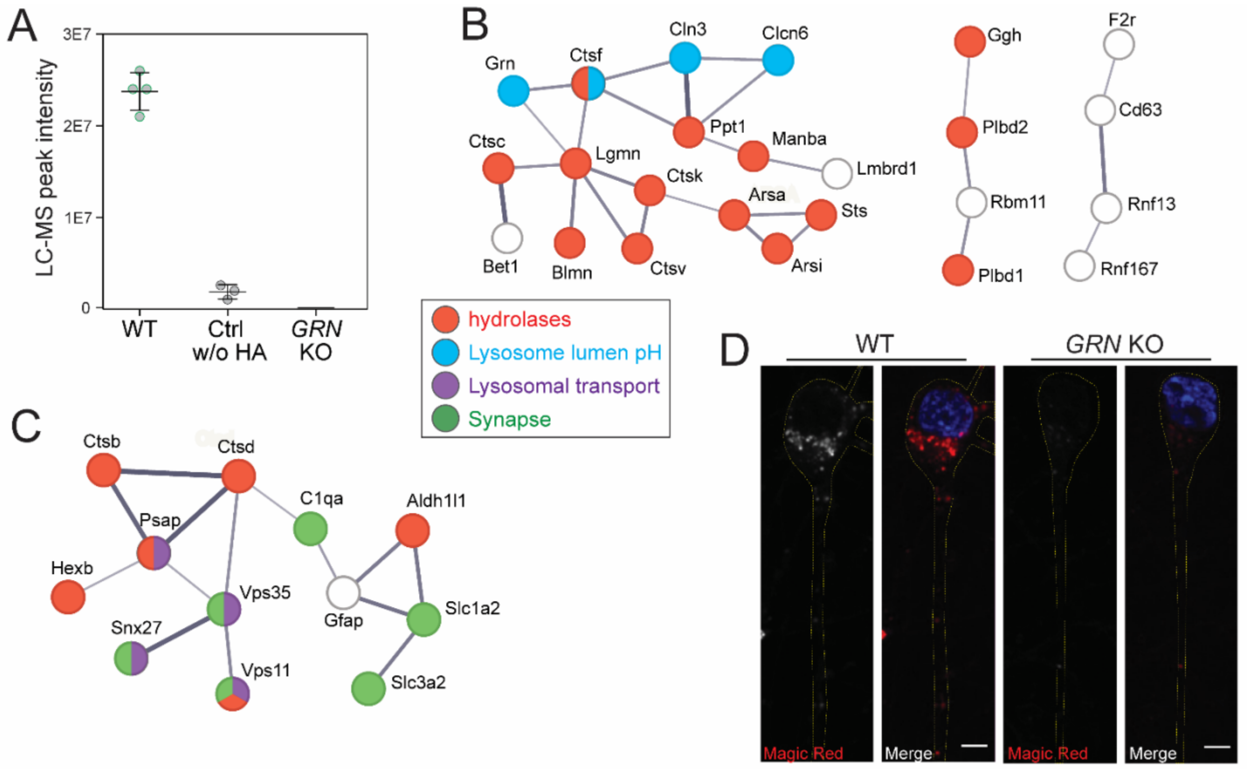


**Supplementary Figure S3. Loss of progranulin results in elevated levels of lysosomal catabolic enzymes and decreased cathepsin B activity in human i$^3$Neurons and mouse brains, related to Figure 3.** (**A**) Confirmation of PGRN enrichment in WT neuronal lysosomes and the loss of PGRN in *GRN* KO i$^3$Neurons using Lyso-IP proteomics. (**B**) Protein network analysis of significantly altered proteins in *GRN* KO vs. WT i$^3$Neurons from Lyso-IP proteomic results. (**C**) Protein network analysis of significantly altered proteins in *GRN*$^{-/-}$ vs. WT mouse brain from Lyso-BAR proteomic results. (**D**) Another replicate of Magic Red assay showed reduced cathepsin B activity in *GRN* KO i$^3$Neurons compared to WT i$^3$Neurons. Scale bar is 10 µm.

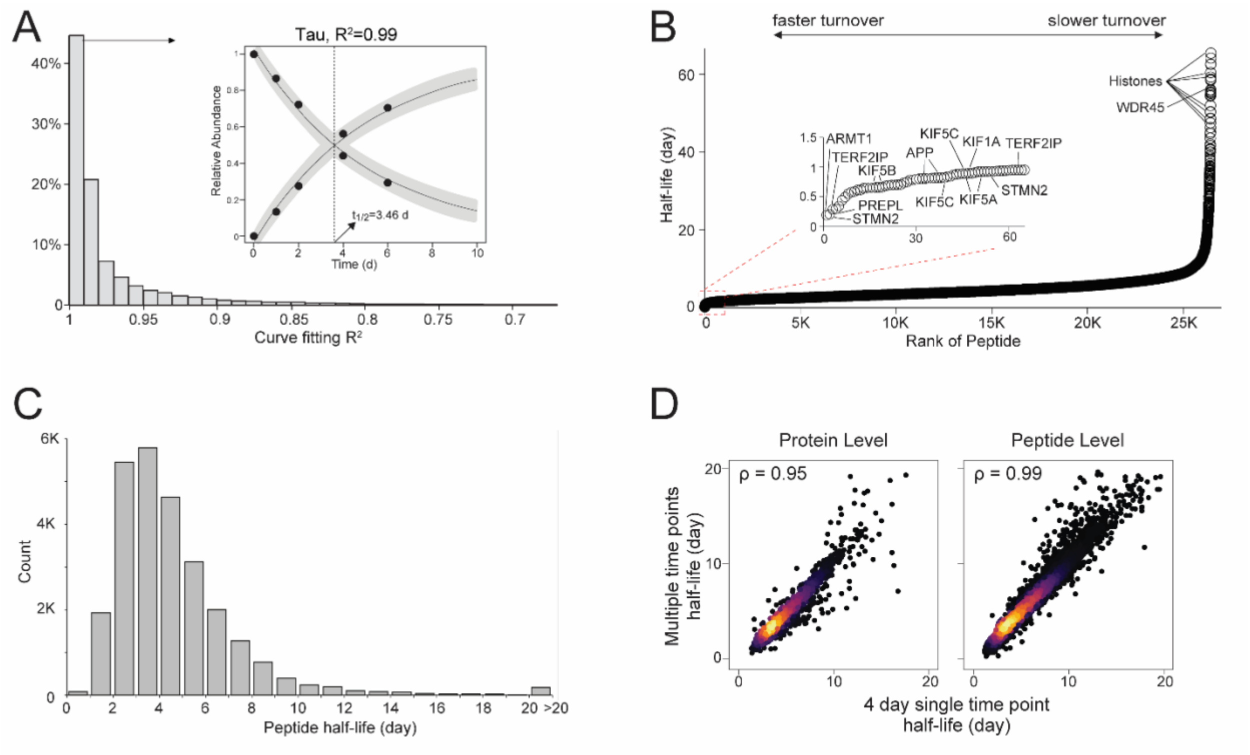


**Supplementary Figure S4. Developing dynamic SILAC proteomics in $i^3$Neurons to measure global protein turnover in cultured human cortical neurons, related to Figure 4.** (**A**) Histogram distribution of $R^2$ in fitting peptide degradation curves to first-order exponential decay. Inlay shows an example of well-fitted Tau peptide degradation and synthesis curves with a half-life of 3.46 day. (**B**) Peptide level half-lives from all unique peptides quantified in WT $i^3$Neurons. (**C**) Histogram distribution of peptide half-lives in consistency with protein level results in Figure 4D. (**D**) Scatter plot showing the correlation of protein/peptide half-lives measured by multiple-time-point method and single-time-point method at day 4 after heavy medium switch.

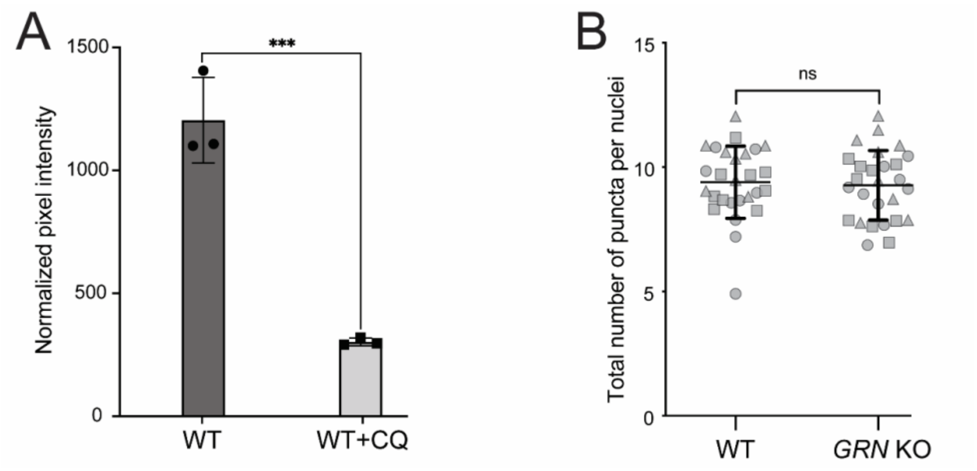


**Supplementary Figure S5. DQ-BSA Red Assay to measure lysosomal degradative function, related to Figure 5.** (**A**) Chloroquine treatment (30µM for 12 hours) significantly reduced the DQ-BSA signals caused impaired lysosomal degradative function in WT i$^3$Neurons (*** denotes *p*-value < 0.001). (**B**) WT and *GRN* KO i$^3$Neurons have similar total number of DQ-BSA fluorescent puncta, indicating similar overall levels of endocytosis and lysosomal biogenesis, related to Figure 5I.

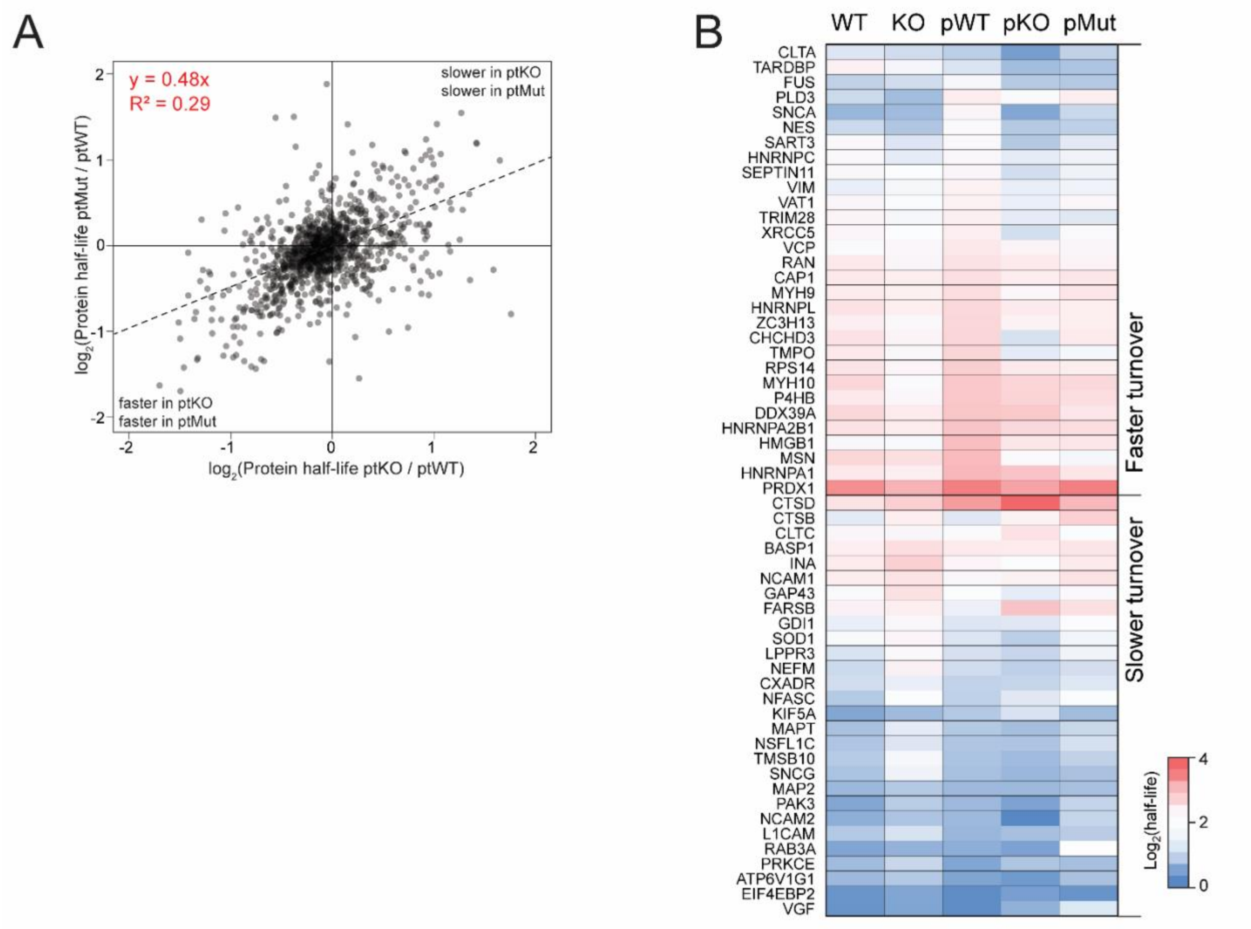


**Supplementary Figure S6**. Protein half-life changes caused by progranulin deficiency in GRN-KO, WT, ptKO, ptMut, and ptWT i$^3$Neurons, related to Figure 6. (**A**) Scatter plot showing potential gene dosage effect of protein half-life changes in ptKO vs. ptWT and ptMutant vs. ptWT i$^3$Neurons. (**B**) Heatmap showing overlapping protein half-lives in WT, *GRN*-KO, ptWT, ptKO, and ptMut i$^3$Neurons. Heatmap colors represent the absolute half-life measurements in days.
